# Threat imminence and estimates of agency dictate proactive and reactive defensive behaviors in active place avoidance

**DOI:** 10.64898/2026.09.11.750960

**Authors:** Alexandre Chambard, Clémence Quittet, Salomé Boyer, Thibaut Vilalta, Jérémie Naudé, Emmanuel Valjent, Federica Bertaso, Antoine Besnard

**Affiliations:** IGF, Univ. Montpellier, CNRS, Inserm, Montpellier, France; INM, Univ. Montpellier, Inserm, Montpellier, France

**Keywords:** behavior, mouse, active place avoidance, defensive behaviors, sex as a biological variable, threat imminence, agency, norepinephrine

## Abstract

Defensive survival behaviors are shaped by both danger attributes and internal physiological states. Well-characterized defensive responses depend not only on whether danger is potential or perceived, but also on its spatial and or temporal proximity. The relationship between danger imminence and defensive behaviors is referred to as the threat imminence continuum, which provides a conceptual framework bridging ecology and behavioral neuroscience. Although this theoretical framework is gaining momentum, another much less explored danger attribute is the extent to which the environment may be controllable. Most paradigms assessing defensive behaviors rely on conditioned stimuli as danger warning-signals in stable environments that hamper the study of environment controllability and its impact on defensive behaviors. To address this question, we repurposed the active place avoidance (APA) paradigm combined with supervised machine learning to characterize multivariate exploratory and defensive behaviors in male and female mice. One important benefit of APA is that it allows the study of active avoidance when the platform is rotating as well as deliberative avoidance when the platform is immobile. As such, APA is very well suited to test the impact of environment controllability on the expression of defensive behaviors. Using this unified preparation, we systematically compared quantitative danger attributes (punishment intensity, behavioral agency) across danger states (absence, presence and omission of punishment) and learning experience (early and late learning stages), while considering sex as a biological variable. Our results demonstrate that defensive behaviors depend on punishment intensity in a sex-dependent manner in APA. Defensive responses can be classified as proactive (risk assessment) or reactive (freezing, escape) in APA. Their response probability depends on behavioral agency, is mutually exclusive and depends on danger spatial proximity. Acute treatment with norepinephrine transporter blocker selectively increases reactive defensive behaviors while sparing proactive defensive behavior. Collectively, these results provide experimental evidence that environmental stability calibrate proactive and reactive defensive behaviors whose neural correlates may be divergent.

- Defensive behaviors depend on punishment intensity in a sex-dependent manner in active place avoidance (APA)
- Environment controllability calibrates proactive and reactive defensive behaviors in APA
- Opposing spatial distribution of proactive and reactive defensive behaviors in APA
- Acute blockade of norepinephrine transporter increases reactive while sparing proactive defensive responses in APA

**Graphical abstract:** Inactive and active place avoidance allow the study of threat imminence as well as estimates of agency. Both task versions are associated with proactive (risk-assessment, avoidance) and reactive (freezing, escape) defensive responses. Enhancing norepinephrine transmission favors reactive defensive responses depending on environment controllability (platform rotation).

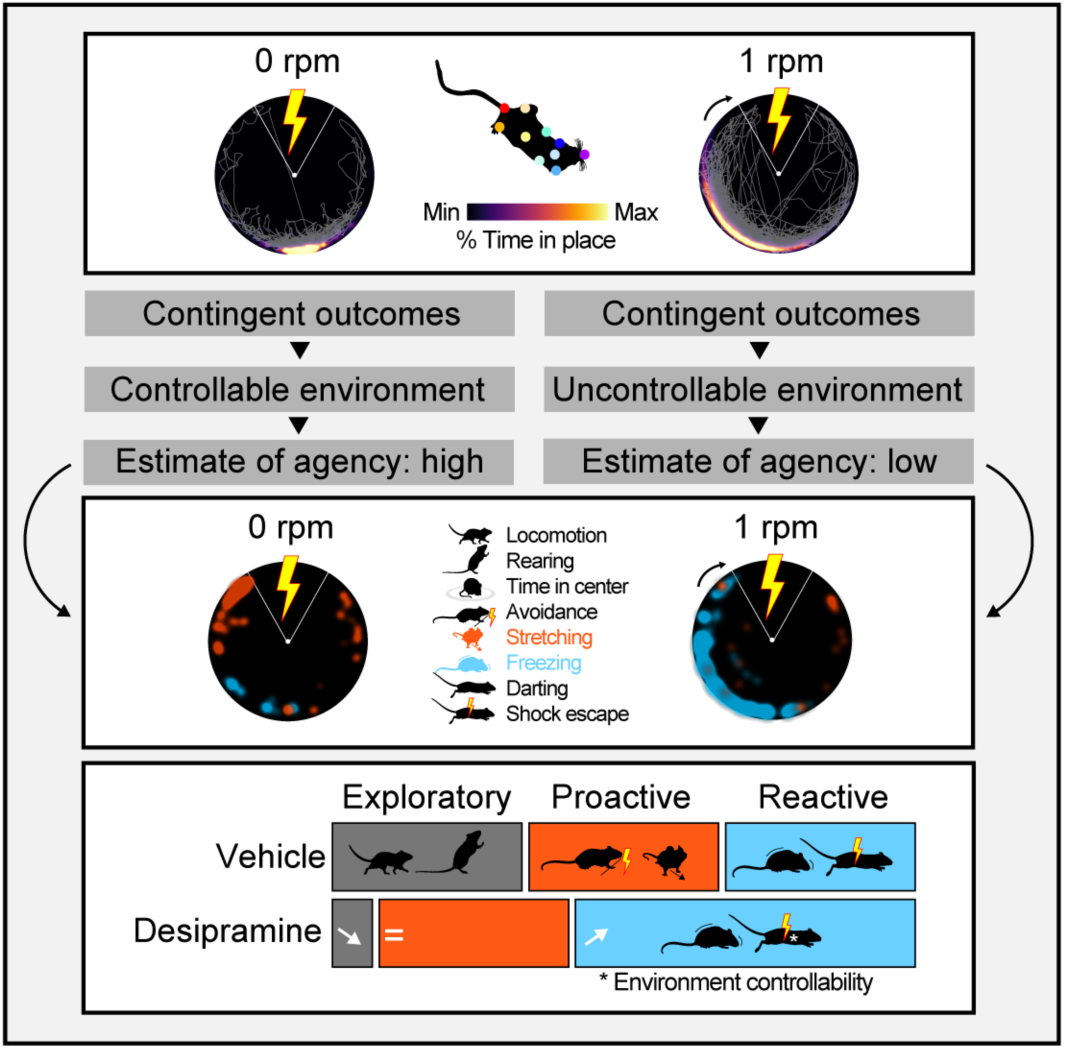

## INTRODUCTION

Defensive survival behaviors are very well conserved across animal species and depend on various factors including body size, physiology and internal states (reproductive status, body condition) [1]. Defensive responses also vary depending on qualitative danger attributes such as whether danger is potential/uncertain or acute/perceived [2]. Quantitative danger attributes also critically shape defensive responses depending on its spatial and/or temporal proximity [1–4]. Distant danger may elicit pre-encounter defensive responses such as avoidance or risk assessment. More imminent danger may elicit post-encounter responses such as freezing behavior. Immediate danger may elicit circa-strike defensive responses such as fight, flight, persistent freezing or tonic immobility [1–4]. The production of heterogeneous defensive responses along the threat imminence continuum provides a conceptual framework bridging ecology [5], behavioral neuroscience in animal models [1–4], and human subjects [6, 7]. It is also highly relevant to psychiatry for the definition of negative valence systems in accordance with NIMH Research Domain Criteria (RDoc) [8], which is further bolstered by empirical evidence linking anxiety disorders and aberrant expression of defensive responses along the continuum [9].

Studies focusing on avoidance of potential danger in rodents rely on conflict-based paradigms that balance fulfilling exploratory drive and self-imposed exposure to potentially harmful situations. For instance, deliberative avoidance of open arms of an elevated plus maze or the exploration of the center area of an open field can be used as proxies of anxiety-like behaviors [10, 11]. In contrast, much of the historical work focusing on avoidance of acute danger has been performed in the two-way shuttle box in which rodents learn to avoid a mild electric footshock by shuttling between two chambers when an auditory cue is presented [12, 13]. The two-way shuttle box requires Pavlovian association of a conditioned stimulus predicting danger followed by avoidance acquired via instrumental conditioning [14]. This assay has allowed gaining insights into the neural circuits mediating goal-directed avoidance [15, 16] and more recently habitual avoidance [17]. Although several well-known issues have limited the interpretation of two-way shuttle box results [14, 18], recent advancements have substantially improved its experimental design thereby circumventing historical interpretational limitations [17, 19].

In recent years, efforts have been dedicated to expand the study of learnt avoidance competing with reward pursue. One such paradigm is the platform-mediated avoidance task that relies on the conflict between pursuing a reward and avoiding a punishment [20]. Platform-mediated avoidance and other similarly inspired paradigms [21, 22] are very well suited to study avoidance that occurs at the expense of other physiological needs [23]. However, it remains unclear to which extent balancing conflicting needs may recapitulate avoiding danger in the pursue of safety [24]. In addition, these paradigms consistently rely on conditioned stimuli as warning signals, and as such also inform the predictive relationship between conditioned stimuli and aversive outcome.

Over the past two decades, active place avoidance (APA) paradigm has been extensively used to study the neural correlate of learning and memory and spatial navigation on a rotating circular arena in rats and mice [25, 26]. APA represents an interesting alternative to traditional models, as it neither relies on explicit cues nor approach-avoidance conflict in order to signal danger or motivate mice to engage with danger. Instead, APA relies on allothetic room-based visual cues to identify and then avoid the punishment zone [27]. The large open-field like environment also allows experimental subjects to display a number of behavioral motifs relevant to assess the competition between exploration and defensive responses. Typical APA training protocol involves a pretraining phase conducted in the absence of punishment and traininig phase in the presence of punishment, which allows longitudinal assessment of behaviors in the absence and presence of acute danger. APA is also well suited to compare behavioral performances in mice trained with a rotating platform as compared to an immobile platform [28], which thus allows to study the impact of environment controllability on defensive behaviors.

Here, we repurposed APA to study multivariate defensive behaviors with markerless pose-estimation analysis [29] to capture multivariate behavioral motifs encompassing exploration (locomotion, rearing, time in center), and defensive responses (avoidance, risk assessment, freezing, darting, footshock escape) [1–4]. These proactive (risk-assessment, avoidance) and reactive (freezing, escape) defensive responses were systematically assessed while varying quantitative danger attributes (punishment intensity, platform rotation) in a longitudinal preparation establishing different danger states (absence, presence and omission of punishment), and experience learning (early and late training stages), while considering sex as a biological variable. Our results demonstrate that defensive behaviors depend on punishment intensity in a sex-dependent manner, and reverse when punishment is omitted during extinction. Both proactive and reactive defensive responses occur in APA. Their respective response probability depends on behavioral agency, is mutually exclusive and depends on danger spatial proximity. Acute treatment with norepinephrine transporter blocker prior to APA or its immobile version selectively increases reactive defensive behaviors while sparing proactive defensive behavior.

Taken together, these results point to APA as a unified preparation capturing multivariate behaviors in a longitudinal manner, indispensable for the study of defensive behaviors along the threat imminence continuum and reflecting estimates of agency. Our results also provide the demonstration that estimates of agency dictate proactive and reactive defensive responses whose neural correlates may be divergent.

## RESULTS

### Active place avoidance and its extinction depend on footshock intensity

Previous work has shown that mice readily learn to avoid a punishment in APA [26]. We reappraised this extensive line of work by varying the footshock intensity in four experimental groups of mice (0, 0.1, 0.2 and 0.3 mA). The protocol consisted in a pretraining session on day 1, training sessions on days 2 and 3 and extinction sessions during which the footshock was omitted on days 4 and 5 (Fig. 1A-B). All groups of mice learnt to avoid the footshock zone, although 0.1 mA footshock yielded a less robust avoidance as compared to 0.2 and 0.3 mA (Fig. 1C-D). A similar relationship between footshock intensity and avoidance was observed during extinction sessions, with 0.3 mA demonstrating the most robust avoidance, which rapidly decayed over time (Fig. 1E). In this assay, we also found that mice rely on distal rather than proximal stationary cues to locate the footshock zone (Fig. S1A-E). Collectively, these results confirm that mice readily learn to avoid a punishment in APA and that response depends on footshock intensity, visual cues present in the environment, and rapidly decays when the foothsock is omitted.

**Figure 1.**
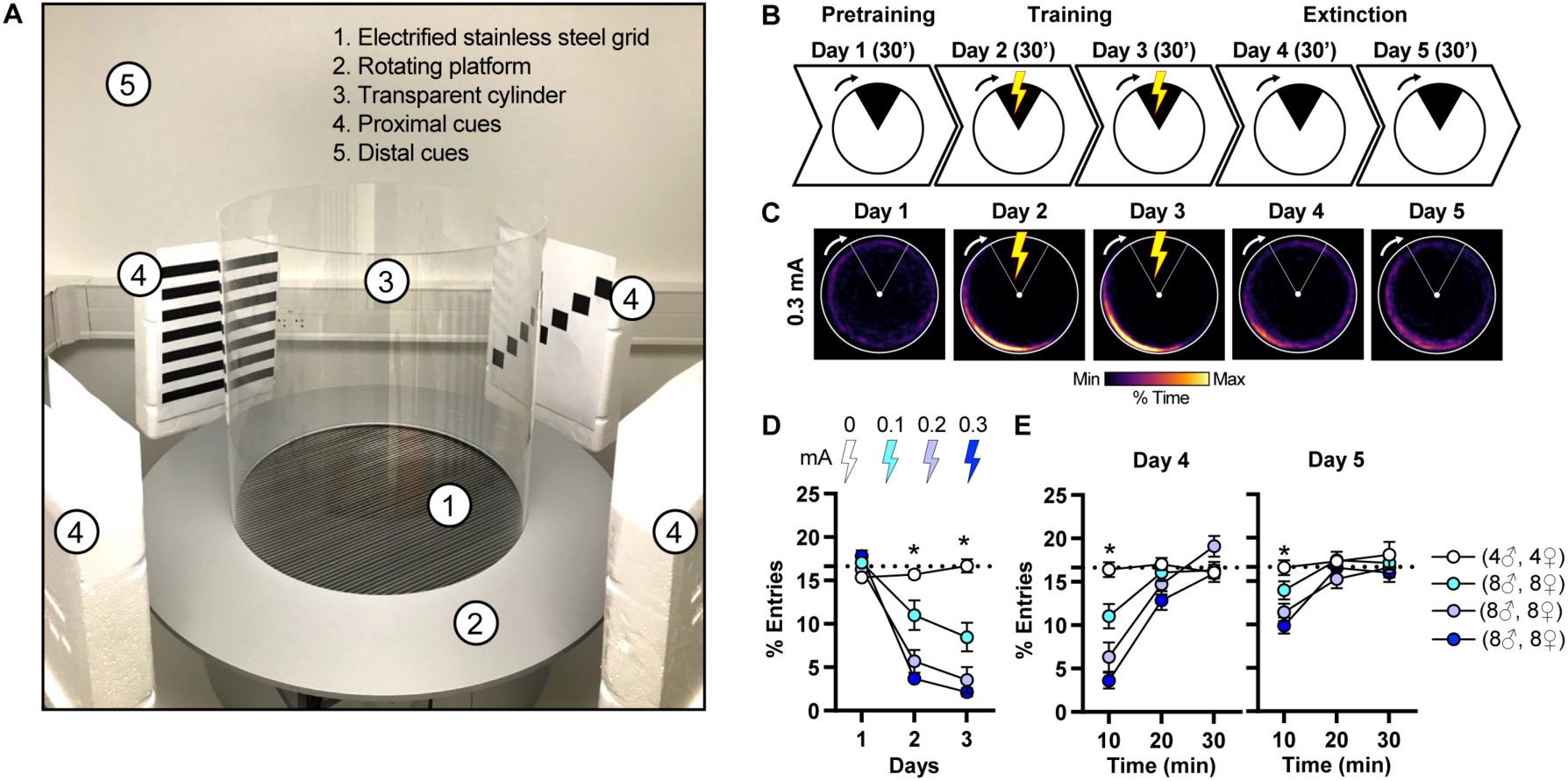
Active place avoidance training and extinction depend on footshock intensity. **A)** APA consists in a stainless steel grid (1) placed on a rotating platform (2). A transparent cylinder (3) prevents mice from escaping. Proximal cues (4) are placed in the vicinity of the platform and distal cues (5) throughout the room. **B)** Active place avoidance and extinction protocol. APA consisted of 1 day of pretraining, 2 days of training and 2 days of extinction. **C)** Time-in-location heat maps for mice trained with 0.3 mA footshocks during each 30 min session. **D)** Mice conditioned with 0.1, 0.2 and 0.3 mA intensity footshocks robustly avoided the shock zone during training. Chance level (dotted line) is set at 16.66% entries. **E)** Mice previously conditioned with higher intensity footshocks demonstrated robust avoidance early on extinction training (Day 4). Data (means ± SEM; n= 8, 16, 16, 16 mice per group) were analyzed using mixed factor two-way ANOVA (repeated measure over time) followed by Tukey multiple comparisons post-hoc test (detailed in Supplementary Table 1), *p < 0.05, 0.3 mA versus 0 mA.

### Markerless pose estimation in APA

The expression of defensive behaviors reflects a repertoire of behavioral motifs that vary depending on threat imminence across pre-encounter, post-encounter and circa strike stages [1–4]. Avoidance and risk assessment have been associated with pre-encounter responses, freezing with post-encounter responses and escape with circa strike responses [1–4]. Avoidance behavior can be readily studied in APA using ‘room frame’ coordinates in which the footshock zone is stationary and the platform rotates clockwise (Fig. 2A). However, key behaviors of interest (e.g., freezing, stretch-attend pose) are based on motion detection and as a result cannot be faithfully measured when the platform is constantly in motion. All subsequent behavioral analyses have thus been performed in ‘arena frame’ coordinates, which reflect ‘room frame’ coordinates with rotation correction. Rotation correction thus enables accurate measurement of locomotion, rearing, percent time in center, stretch-attend pose (referred to as stretching), freezing, darting and escape vigor (Fig. 2A). To capture these behavioral features, we combined APA with DeepLabCut markerless pose estimation analysis [29]. We used 10 different tracklets to monitor the head, chest and back centroids of mice (Fig. 2B-C) that could then be visualized in ‘room frame’ and ‘arena frame’ (Fig. 2D). We then systematically compared DLC behavioral measurements with a commercial videotrack or two blinded experimenters on mice that were trained with 0.1, 0.2 and 0.3 mA in the initial APA experiment (Fig. S2). DLC reached commercial and manual scoring performances for horizontal locomotion (Fig. S2A), percent time spent in center (Fig. S2B), supported rearing (Fig. S2C), and freezing (Fig. S2D-G). DLC measurements captured stretch-attend pose referred to as stretching (Fig. S2H), which is an ethologically relevant behavior that is highly sensitive to anxiolytic treatments [30, 31] and has been largely neglected, as it is difficult to consistently monitor with the naked eye [32]. DLC measurements also allowed to capture footshock escape vigor (Fig. S2I) and darting events (Fig. S2J), which display the characteristics of learned defensive behaviors [33]. We then applied this analysis pipeline to measure all behaviors in mice trained with different footshock intensities in APA (Fig. 1) across 5 sessions (Fig. 2E-K). Mice trained with footshocks showed lower levels of locomotion, rearing and time in center compared to control mice (Fig. 2E-G). They also showed greater levels of stretching and freezing (Fig. 2H-I). There was no difference in the number of darting events (Fig. 2J) and escape vigor was stable across training sessions (Fig. 2K). Importantly, all behavioral measures that were altered during training reversed to control levels during extinction except for the percent time spent in center, which remained lower in mice previously trained with 0.3 mA (Fig. 2G). Altogether, these results indicate that supervised pose estimation analysis captures the dynamic competition between exploratory behaviors and defensive behaviors during APA training, which can reverse to baseline conditions during extinction (Fig. 2L). Notably, we found that stretch-attend pose, freezing and footshock escape vigor are all differentially impacted by varying footshock intensities (Fig. 2L-M). While risk assessment (stretching) was observed across all footshock intensities, freezing was only observed with higher footshock intensities. In the same vein, escape behavior was more pronounced with higher footshock intensities (Fig. 2L-M). The expression of all defensive behaviors decreased during extinction training allowing the re-emergence of exploratory behaviors. Collectively, these results indicate that APA is well suited to study distinct danger states ranging from deliberative exploration in the absence of perceived danger (pretraining), acute danger (training), and immediate danger (exposure to footshock).

**Figure 2.**
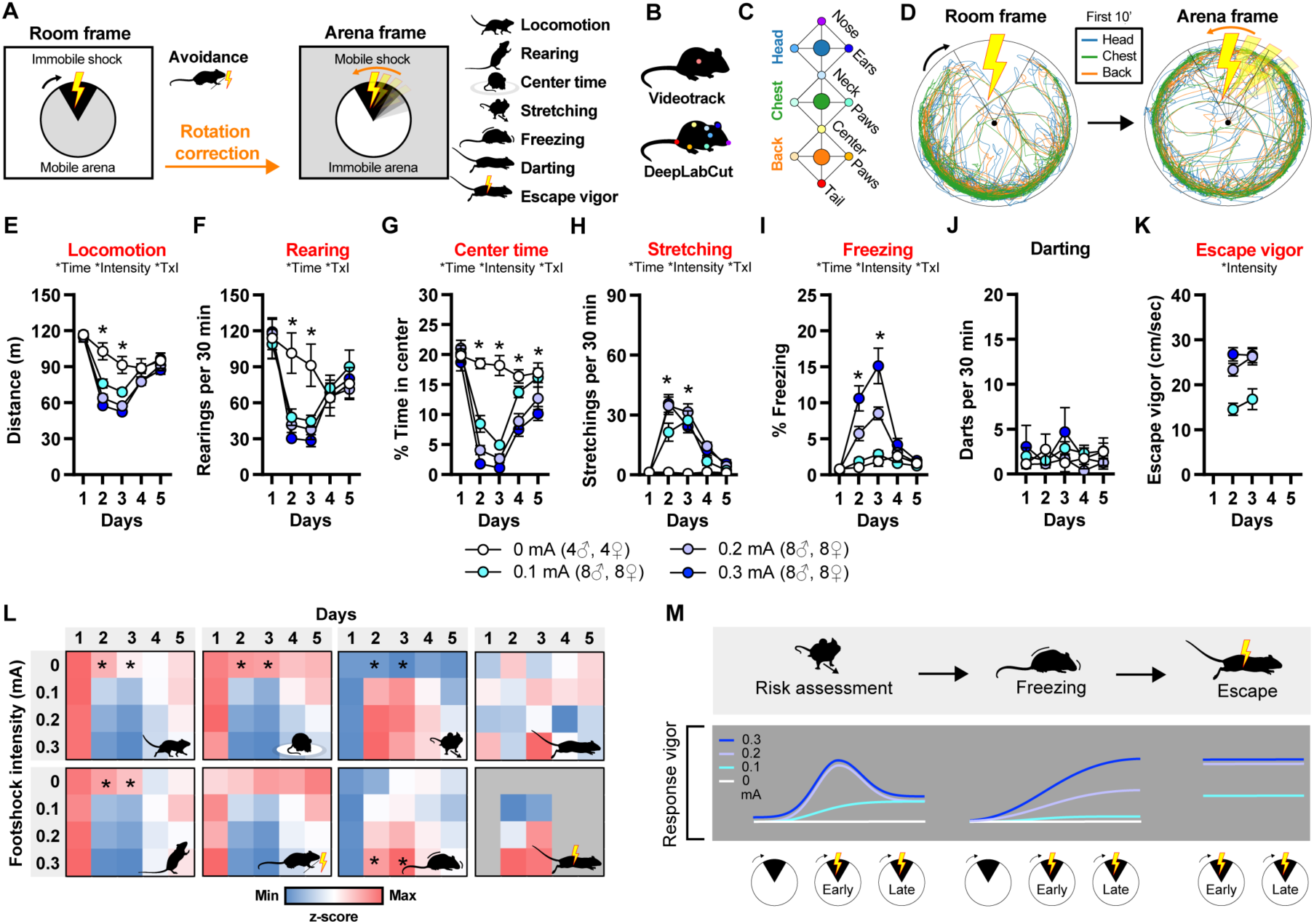
Pose estimation analysis unveils multivariate exploratory and defensive behaviors in APA. **A)** Schematic illustrating the expression of mouse position in two reference frames. Converting mouse coordinates from “Room frame” to “Arena frame” allows the assessment of multivariate behaviors. **B)** Analysis of avoidance behavior was performed with a videotracking software and other behaviors with DeepLabCut (DLC) pose estimation toolbox. **C)** Pose estimation was performed by tracking 10 individual tracklets allowing precise reconstruction of 3 centroids, namely, head, chest and back. **D)** Representative paths of each centroid plotted in “Room frame” and “Arena frame” from one mouse in the first 10 min of a training session. **E-K)** Pose estimation analyses of locomotion (E), rearing (F), percent time in center (G), stretching (H), freezing (I), darting (J) and escape vigor (K) across the 5 days protocol in all groups of mice. Red titles denote significant differences for clarity. Data (means ± SEM; n= 8, 16, 16, 16, 8, 8 mice per group) were analyzed using Mixed-effect model followed by Tukey multiple comparisons post-hoc test (detailed in Supplementary Table 1), *p < 0.05, 0.3 mA versus 0 mA. Main effect of time or intensity as well as interaction (TxI) are indicated for each comparison. **L)** Raster plots and summary statistics depicting behavioral motifs usage expressed as z-scores on each day (1-5) across different footshock intensites (0 to 0.3 mA). **M)** Threat intensity differentially impacts the vigor of risk assessment and freezing, as well as circa-strike responses (escape vigor).

### Sex-specific differences fade with repeated training and depend on footshock intensity in APA

The expression of defensive behaviors varies depending on a number of factors including biological sex [34]. We thus revisited all behaviors in mice trained with different footshock intensities in APA on day 3 (Fig. 2) taking into consideration sex as biological variable and different footshock intensities (Fig. 3A). We used all 8 behavioral features (Fig. 3B) to train a random forest classifier to predict experimental group membership (Fig. 3C). Overall classification exceeded a shuffled distribution, and this effect was mostly driven by the accurate classification of footshock intensity (Fig. 3D) rather than sex (Fig. 3E). Although the effect of footshock intensity may outweigh the effect of sex, the confusion matrix indicated accurate classification of sex at 0.1 mA (Fig. 3C). We thus assessed the effect at various footshock intensities separately (Fig. 3F, S3A-X). Discrete sex-specific differences emerged at different footshock intensities whereby female mice showed greater levels of risk assessment and escape vigor at low footshock intensities (0.1 mA) and greater freezing at higher footshock intensities (0.2-0.3 mA) (Fig. 3F-I, S3A-X). Importantly, these differences were more prominent early on training (day 2) and seemingly disappeared later on training (day 3). We thus tested whether all behaviors allow accurate classification of sex across days (pretraining, training and extinction) in mice trained with different footshock intensities (Fig. 3J). We trained random forest classifiers to predict sex from behaviors (Fig. 3K). Sex classification exceeded a shuffled distribution on the first training session (day 2) at all footstock intensities (Fig. 3L). These results demonstrate that multivariate behaviors in APA temporarily contain predictive information resolving sex across different footshock intensities. The observation that APA consistently differs across sexes naturally prompted us to evaluate if these differences reflect hormonal variations across the estrous cycle. We trained an additional cohort of female mice in an abbreviated version of APA (2 days) (Fig. 3M). We then performed vaginal cytology in order to establish the estrous cycle stage upon completion of APA training (Fig. 3N) and sorted mice in three subpopulations in either low hormonal state (estrus and diestrus) or high hormonal state (proestrus) [34]. Interestingly, variations in the estrous cycle did not influence any of the exploratory or defensive behaviors in APA training (Fig. 3O-P, S3Y-AE). Finally, a random forest classifier trained on behavioral data from day 2 failed to successfully classify mice belonging to each estrous stage (Fig. 3Q-R). These results do not support the a contribution of the estrous cycle in the transient sex-differences observed early on in APA (day 2).

**Figure 3.**
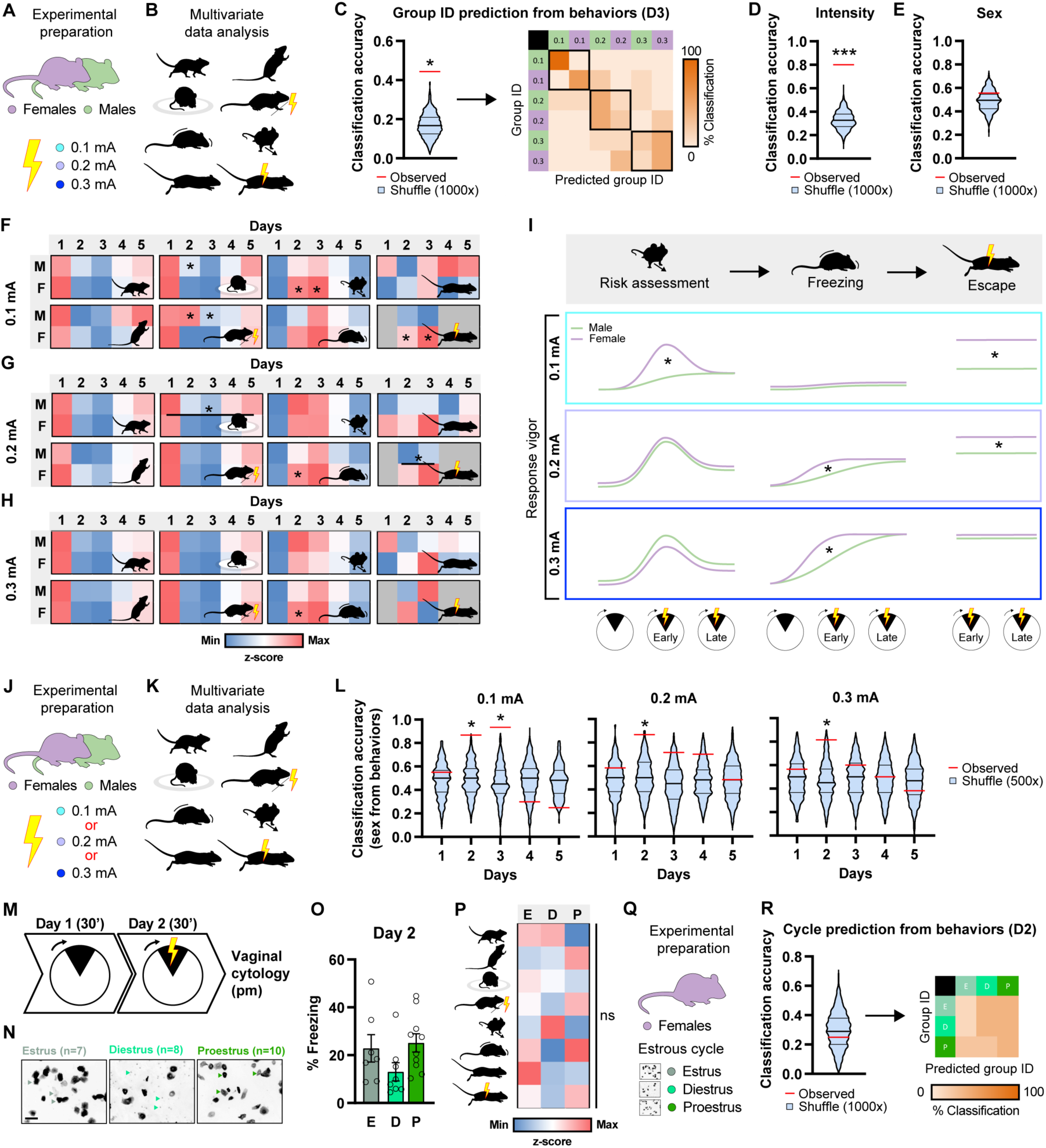
Female mice display greater defensive behaviors along the threat imminence continuum in APA. **A-B)** Schematic illustrating the experimental conditions on day 3 taking into consideration both sexes and 3 footshock intensities (A) across 8 behavioral measurements (B). **C)** All 8 behaviors were used to train a random forest classifier in order to predict group identity from behavioral measures. The confusion matrix depicts accurate classification of mice from both sexes and all footshock intensities. Data (n= 8 mice per group, 48 mice total) were analyzed using one-tailed non-parametric permutation test (detailed in Supplementary Table 1), *p < 0.05, observed versus shuffle. **D-E)** The same analysis was conducted on either footshock intensity (D) or sex (E). **F-H)** Raster plots and summary statistics depicting behavioral motifs usage expressed as z-scores on each day (1-5) across sexes for mice trained with 0.1 mA (F), 0.2 mA (G) and 0.3 mA (H). **I)** The vigor of risk assessment and freezing as well as circa-strike responses vary depending on sex and footshock intensities. **J-K)** Schematic illustrating the experimental conditions taking into consideration both sexes and 3 footshock intensities (J) across 8 behavioral measurements (K). **L)** All 8 behaviors were used to train random forest classifiers in order to predict sex from behavioral measures on each day for all footshock intensities. Data (n= 16 mice per group) were analyzed using one-tailed non-parametric permutation test (detailed in Supplementary Table 1), *p < 0.05, observed versus shuffle. **M)** Female mice were trained in a two-day APA protocol. **N)** At the end of day 2, the estrous cycle was monitored and mice were sorted based on the relative density of cornified cells (gray arrowheads, estrus), leukocytes (cyan arrowheads, metestrus and diestrus) and nucleated cells (green arrowheads, proestrus). **O)** Freezing behavior is not different across mice at different stages of the estrous cycle. **P)** Raster plot depicting behavioral motifs usage expressed as z-scores on day 2 across mice in estrus, diestrus and proestrus. **Q)** Schematic illustrating the experimental conditions on day 2 taking into consideration the three different stages of the estrous cycle. **R)** All 8 behaviors were used to train a random forest classifier that failed to accurately classify group identity from behavioral measures. Data (n= 7 mice per group, 21 mice total) were analyzed using one-tailed non-parametric permutation test (detailed in Supplementary Table 1), *p < 0.05, observed versus shuffle.

### Proactive and reactive defensive behaviors are mutually exclusive and depend on environment controllability

Our observation that risk assessment and freezing emerge early and late in APA, respectively, suggested that mice re-appraise their perception of danger as they refine knowledge about danger attributes. We wondered whether this opposing pattern was observed as a result of limited environment controllability owing to constant platform rotation. To test this hypothesis, we took advantage of one critical feature of APA that is the possibility to switch on or off platform rotation, as mice readily avoid an air puff on a stationary platform [28]. We thus questioned to which extent platform rotation impacts the balance between risk assessment and freezing behaviors by training an additional group of mice on a stationary platform using 0.3 mA footshocks (Fig. 4A-B). Mice trained in the inactive version learned and extinguished footshock avoidance as efficiently as mice trained in the active version (Fig. 4C-D). We then deployed the same DLC analysis pipeline in the inactive version of the task, without applying rotation correction (Fig. S4A-J). When comparing other behaviors, mice were overall less mobile in the inactive version (Fig. 4E) and no major differences were observed on rearing, time in center and footshock escape vigor during training sessions (Fig. 4F, G, K). However, mice trained in the inactive version showed substantially less freezing and more stretching during training sessions (Fig. 4 H-I) and more darting events during extinction (Fig. 4J). These results demonstrate that although mice successfully learn to avoid the footshock zone regardless of platform rotation, active and inactive place avoidance elicit contingency-specific defensive behaviors (Fig. 4L). Specifically, mice engage substantially more in risk assessment in the inactive version of the task as compared to mice trained in the active version (Fig. 4M). Importantly, expression of risk assessment and freezing did not differ across sexes in the inactive version of the task (Fig. 4N, Fig. S4O). However, a random forest classifier trained on behavioral data across days successfully predicted sex during the training sessions (Fig. 4O-R). These results likely reflect differences in exploratory behaviors (locomotion, time in center) across male and female mice in the inactive version of the task (Fig. S4K-M). Lastly, we compared the spatial distribution of stretching and freezing events on day 3 in the inactive and active versions of the task (Fig. 4S). Both stretching and freezing events were expressed in a symmetrical pattern in the inactive version, and this pattern was skewed towards the footshock zone in the active version of the task (Fig. 4T-U). Interestingly, freezing and stretching occurred in a mutually exclusive manner, whereby stretching was most observed in the vicinity of the footshock zone and freezing away from the footshock zone (Fig. 4T-U). Altogether, these results demonstrate that manipulating platform dynamics in APA offers the opportunity to investigate the transition from proactive (risk assessment) to reactive (freezing) defensive behaviors and their putative neural correlates.

**Figure 4.**
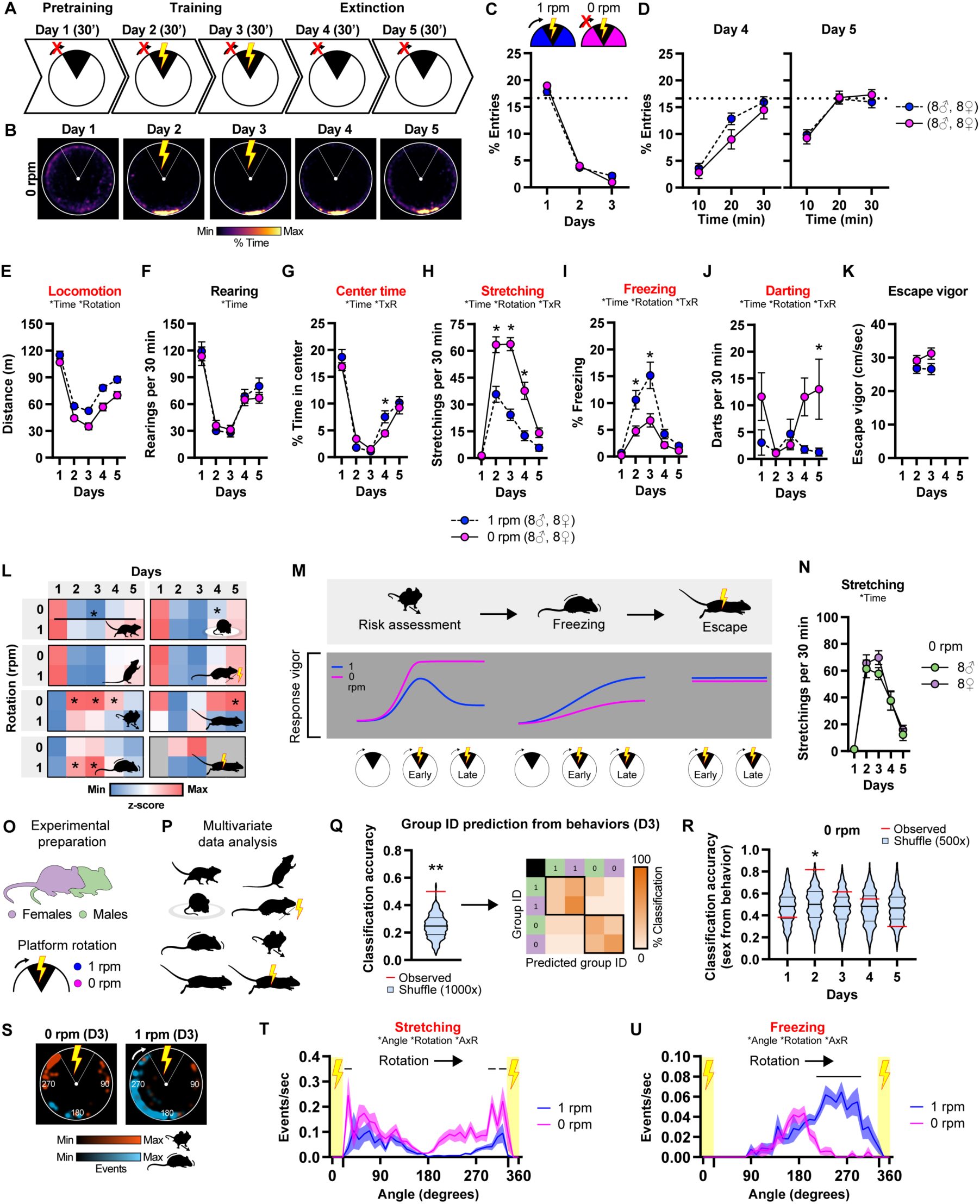
Estimates of agency as a critical regulator of defensive behaviors in APA. **A)** Schematic illustrating inactive place avoidance during the pretraining, training and extinction phases. **B)** Time-in-location heat maps for mice trained with 0.3 mA footshocks on a stationary platform. **C-D)** No difference in the acquisition (C) and extinction (D) of active (1 rpm) and inactive (0 rpm) place avoidance. Mice undergoing extinction in the active version were the same as the group presented in Fig. 1 and were only used for direct comparison. Data (means ± SEM; n= 16, 16 mice per group) were analyzed using mixed factor two-way ANOVA (repeated measure over time) followed by Bonferroni multiple comparisons post-hoc test (detailed in Supplementary Table 1), *p < 0.05, cues versus no cues. **E-K)** Pose estimation analyses of locomotion (E), rearing (F), percent time in center (G), stretching (H), freezing (I), darting (J) and escape vigor (K) across the 5 day protocol in both groups of mice. Red titles denote significant differences for clarity. Data (means ± SEM; n= 16 mice per group) were analyzed using mixed-effect model followed by Bonferroni multiple comparisons post-hoc test (detailed in Supplementary Table 1), *p < 0.05, 1 rpm versus 0 rpm. Main effect of time or rotation as well as interaction (TxR) are indicated for each comparison. **L)** Raster plots and summary statistics depicting behavioral motifs usage expressed as z-scores on each day (1-5) across mice trained with 0 and 1 rpm. **M)** The vigor of risk assessment and freezing vary depending on platform rotation. **N)** Stretching behavior is indistinguishable between male and female mice across the 5 day protocol in on the inactive platform. **O-P)** Schematic illustrating the experimental conditions on day 3 taking into consideration both sexes and platform rotation speed (O) across 8 behavioral measurements (P). **Q)** All 8 behaviors were used to train a random forest classifier in order to predict group identity from behavioral measures. The confusion matrix depicts accurate classification of mice from both sexes and rotation speeds. Data (n= 8 mice per group, 32 mice total) were analyzed using one-tailed non-parametric permutation test (detailed in Supplementary Table 1), *p < 0.05, observed versus shuffle. **R)** All 8 behaviors were used to train random forest classifiers in order to predict sex from behavioral measures on each day on the immobile platform. Data (n= 16 mice per group) were analyzed using one-tailed non-parametric permutation test (detailed in Supplementary Table 1), *p < 0.05, observed versus shuffle. **S) H**eat maps for stretching events (red) and freezing events (blue) for mice trained with 0.3 mA footshocks on a stationary platform (0 rpm) or mobile platform (1 rpm) on day 3. **T-U) E-K)** Spatial distribution of stretching (T) and freezing (U) events frequency in both groups of mice. Data (means ± SEM; n= 16 mice per group) were analyzed using mixed-effect model followed by Šídák multiple comparisons post-hoc test (detailed in Supplementary Table 1), p < 0.05, 1 rpm versus 0 rpm.

### Enhancing norepinephrine transmission promotes reactive defensive behaviors in APA

Our observation that risk assessment and freezing occur in a mutually exclusive manner in APA suggested that the underlying mechanism involves competitive or inhibitory interactions. Previous work points to heightened noradrenergic tone following an aversive event as a critical modulator of aversive learning processes under different levels of behavioral arousal [35]. We thus used the selective norepinephrine transporter inhibitor desipramine in order to test whether accumulation of extracellular norepinephrine levels may differentially impact proactive and reactive defensive behaviors in a platform rotation-dependent manner. We first performed control experiments to evaluate whether an acute systemic injection with desipramine (20 mg/kg, i.p.) alters spontaneous exploratory behaviors in an open field (Fig. S5A). We found that desipramine enhances behavioral habituation (Fig. S5B-F) by decreasing locomotor behavior and increasing immobility in a time-dependent manner (Fig. S5C, S5E), without interfering with other spontaneous exploratory or defensive behaviors (Fig. S5G-J). We then performed acute injections with desipramine prior to contextual fear conditioning (CFC) and retrieval and found no difference in freezing behavior as well as escape behavior elicited by 0.3 mA electric footshocks (Fig. S5L-M). However, acute desipramine enhanced freezing behavior in a brief 3 min retrieval session taking place 24 hour after conditioning, thus suggesting that desipramine could promote the stabilization and/or retrieval of contextual fear conditioning (Fig. S5N). Collectively, these results indicate that acute systemic treatment with desipramine favors behavioral habituation in a novel environment as well as promotes the expression of reactive defensive responses in a context previously paired with a punishment. Since desipramine enhances the expression of freezing behavior, we next tested whether acute systemic treatment with desipramine could interfere with active place avoidance and related defensive behaviors in a platform rotation dependent manner (Fig. 5A-D). Acute systemic injection with desipramine decreased exploratory behaviors (Fig. 5E-F) without interfering with center avoidance or entries into the footshock zone (Fig. 5G-H) as well as risk assessment (Fig. 5I). However, desipramine potently enhanced freezing behavior in both versions of the task (Fig. 5J), which was accompanied by an increased in darting and escape vigor only when the platform was rotating (Fig. 5K-L). Spatial distribution of stretching, freezing and darting events in the active version of the task (day 3) revealed an overall increase in freezing behavior that matched the spatial distribution of control mice (Fig. 5M). Desipramine preferentially decreased exploratory behaviors in male mice (Fig. S5O-P) but increased reactive defensive behaviors in both sexes (Fig. S5S). Altogether, these results demonstrate that acute systemic enhancement of noradrenergic tone selectively promotes reactive defensive behaviors (freezing) while sparing proactive defensive behaviors (risk assessment) (Fig. 5N). These effects are much more pronounced in situations of low environment controllability when the platform is active, leading to an enhancement of darting and escape vigor (Fig. 5N).

**Figure 5.**
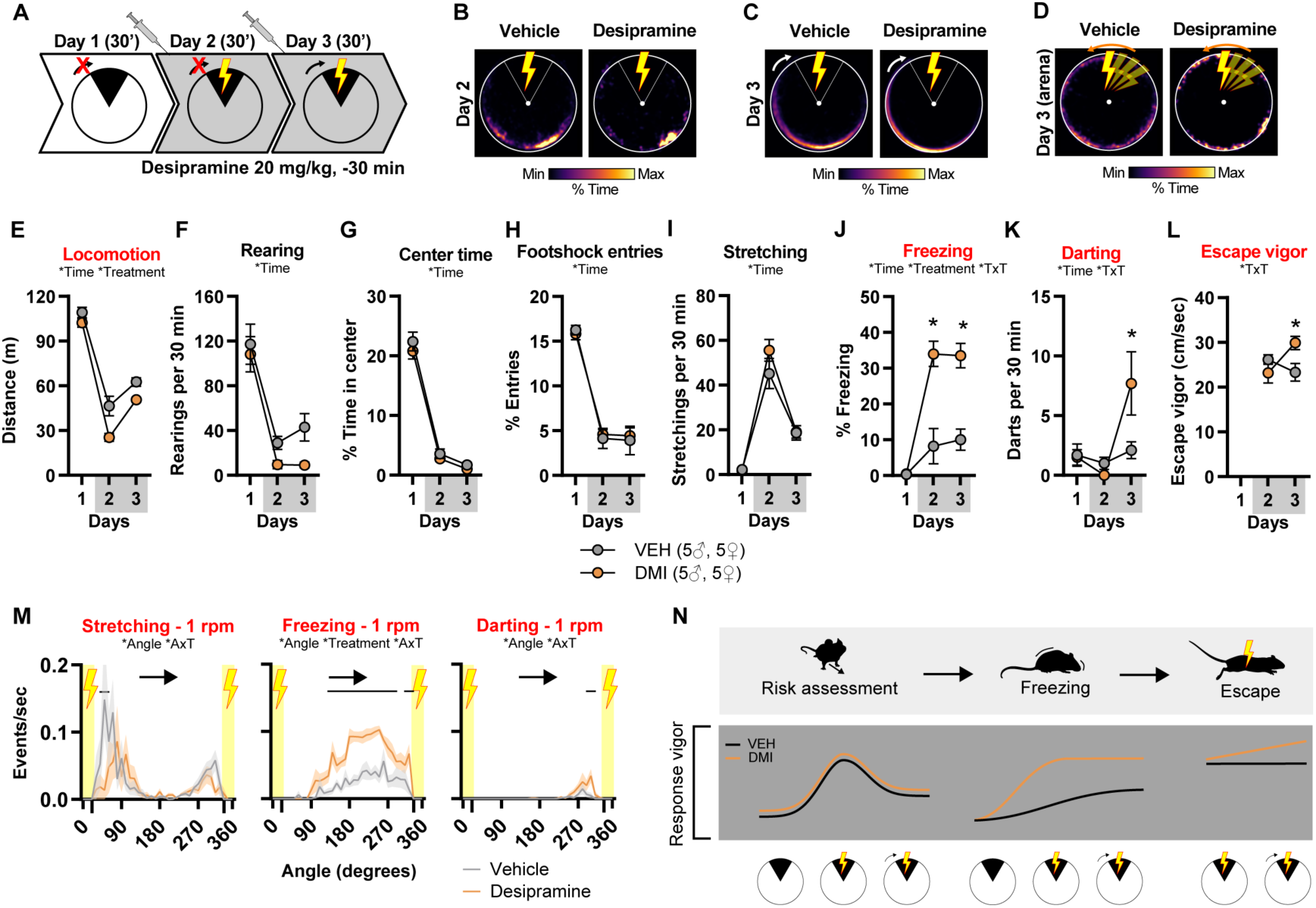
Enhancing norepinephrine transmission promotes reactive defensive responses in APA. **A)** Schematic illustrating acute injections with desipramine prior to inactive followed by active place avoidance. **B-D)** Time-in-location heat maps for mice trained with desipramine or its vehicle on the stationary platform (B) and active platform in room frame (C) or arena frame (D). **E-L)** Pose estimation analyses of locomotion (E), rearing (F), percent time in center (G), footshock entries (H), stretching (I), freezing (J), darting (K) and escape vigor (L) across the 3 days protocol in both groups of mice. Red titles denote significant differences for clarity. Data (means ± SEM; n= 10 mice per group) were analyzed using mixed factor two-way ANOVA (repeated measure over time) followed by Bonferroni multiple comparisons post-hoc test (detailed in Supplementary Table 1), *p < 0.05, desipramine versus vehicle. Main effect of time or treatment as well as interaction (TxT) are indicated for each comparison. **M)** Spatial distribution of stretching, freezing and darting events frequency in both groups of mice. Data (means ± SEM; n= 10 mice per group) were analyzed using mixed-effect model followed by Šídák multiple comparisons post-hoc test (detailed in Supplementary Table 1), p < 0.05, desipramine versus vehicle. Main effect of angle or treatment as well as interaction (AxT) are indicated for each comparison.

## DISCUSSION

The active place avoidance (APA) assay has been introduced two decades ago and has been used to study the neural correlate of learning and memory as well as spatial navigation in rats and mice [25, 26]. In the present study, we repurposed APA to capture multivariate defensive behaviors across danger states. APA involves different learning phases in the absence or presence of a punishment, offering unique opportunity to directly compare how the absence (pretraining) or presence (training) of danger, as well as danger omission (extinction), impact multivariate defensive behaviors in a unified preparation. In the present work, we first confirmed that APA can readily be established with low intensity footshocks (0.2-0.3 mA)[36]. We also confirmed that once established, the omission of the footshock leads to a prompt extinction of the initial avoidance response [36]. Our results also provide demonstration that distal stationary visual cues are indispensable for locating the footshock zone during the extinction phase. In other words, mice do not rely on other discrete olfactory or auditory cues that could potentially confound the interpretation of APA. This latter observation is in line with previous work showing that omitting visual cues prevents mice from locating the footshock zone during the acquisition phase of APA [27].

Recent initiatives have led to the recognition of treating sex as a biological variable [37], which is especially relevant owing to sex differences in anxiety and mood disorders [34]. Previous work has demonstrated sex-specific differences in rat active place avoidance [25]. Here, we show discrete sex-specific differences depending on footshock intensity. At 0.1 mA, male mice show less avoidance and more exploratory behavior, and at 0.2-0.3 mA female mice show greater freezing behavior. These results could reflect a difference in footshock sensitivity or behavioral adaptation to footshock, which has previously been reported in auditory fear conditioning [38]. It should be noted, however, that sex-specific differences were most evident early on training. Accordingly, while multivariate analysis reliably decoded sex above chance, this should be interpreted as evidence that behavioral structure carries sex-related information rather than as evidence of pervasive sexual dimorphism in defensive responding. Nevertheless, our results also indicate that male and female mice avoid just as well using 0.2-0.3 mA, in contrast with recent findings using the two-way shuttle box in which female mice outperformed males at different footshock intensities ranging from 0.1 to 0.3 mA footshocks [39]. Importantly, sex-specific differences using 0.3 mA were mostly noticeable in the exploratory behaviors detected in the inactive version of the task. These results thus suggest that agency, or lack thereof, that male and female mice exert over footshock may be critical for the expression of sex-specific differences in this task.

APA has been traditionally measured with absolute number of entries into the footshock zone [36]. Here we complemented this measurement by performing markerless pose estimation analysis with DeepLabCut [29]. Based on the taxonomy of defensive behaviors across the threat imminence continuum [1–4], we chose to focus on specific behavioral motifs that encompass exploratory behaviors (locomotion, rearing, time spent in center), pre-encounter behaviors that occur during risk assessment (stretch-attend posture), post-encounter behaviors (freezing, darting) and circa-strike behaviors (footshock escape vigor). The ability to analyze these multivariate defensive behaviors together with exploratory behaviors in APA departs from the literature, in that most avoidance protocols focus on post-encounter and circa strike responses [18]. The inclusion of stretch-attend posture [32] is an important step towards understanding risk assessment that constitutes bona fide pre-encounter behavior which is becoming increasingly taken into consideration [40] as well as other behaviors such as scanning behavior [41]. In accordance with the threat imminence continuum framework, our results suggest that exploratory behaviors are much more prominent during the pretraining phase in the absence of danger. Pre-encounter behaviors rapidly increase early on during training and slightly decay at later stages of training, which could reflect the emergence of competitive post-encounter behaviors. Circa-strike behaviors are stable across training sessions. Importantly, most pre- and post-encounter behaviors quickly decayed whenever the footshock was omitted during extinction sessions, which clearly suggests the associative nature of these behaviors. Finally, our analysis pipeline also included darting behavior [33], which we found expressed at low levels in the active version of the task. Altogether, our results clearly demonstrate that APA elicits a range of defensive responses encompassing pre- and post-encounter behaviors as well as circa-strike responses, all pertaining to the threat imminence continuum.

One key feature of APA is that mice use spatial cues to identify the footshock zone and platform rotation fosters engagement with danger in the absence of conditioned stimuli. This is in stark contrast with other behavioral assays such as the approach-avoidance task or the two-way shuttle box, for which the footshock is signaled with a conditioned stimulus [14]. In the latter task, high levels of freezing are often reported which may interfere with active avoidance, although recent work has suggested ways to circumvent this limitation [19]. In APA, learning the localization of the footshock zone occurs via incidental association processes. This specific feature may explain at least in part why such low freezing levels are observed in this task. An alternative interpretation (not at odds with the former) is that APA takes place in an open arena for which the animals exert large control over the onset of the punishment, as opposed to the two-way shuttle box, which is dictated by the experimenter. Controllability of an aversive unconditioned stimulus is crucial in determining subsequent avoidance performance [42, 43]. The two assays may thus differ in their respective degree of controllability, whereby APA may favor avoidance behavior at the expense of freezing behavior owing to low threat imminence. Although APA and two-way shuttle box differ greatly with regard to freezing behavior, we still observed a minority of mice (approximately 10%) that failed to avoid the footshock zone across both training sessions. These so-called non-avoider mice have consistently been reported in the two-way shuttle box [19].

In light of the spatial nature of APA, we thought to consider the expression of defensive behaviors in a two-dimensional space relative to the footshock zone. One important finding was that freezing occurred away from the shock zone while risk assessment (stretching) occurred in the vicinity of the footshock zone. While these results clearly supported the notion that defensive behaviors may reflect the spatiotemporal proximity of danger, they failed to explain how mice may toggle between pre- and post-encounter behaviors when danger has clearly been perceived and is constant across training sessions.

To tackle this important question, we took advantage of one important feature of APA that is the opportunity to modulate the level of environment controllability in the presence of acute danger. To do so, we directly compared defensive behaviors elicited in the inactive and active versions of the task. These results clearly demonstrated that the expression of defensive behaviors does not just reflect the threat imminence continuum but also the level of control that mice exert over the environment. When the environment is highly controllable (inactive platform), mice preferentially engage in risk assessment. When the environment becomes less controllable (active platform), mice preferentially engage in freezing behavior. These results are in accordance with the notion that estimates of agency calibrate an individual’s response along a behavioral continuum ranging from proactive to reactive responses [44]. In situations affording little opportunity for control, an individual can rely on reactive strategies to cope with behavioral challenges [44]. In this framework, freezing, darting and footshock escape are very well-suited reactive responses in APA. Conversely, proactive behavior is characterized by the tendency to explore and discover the structure of reinforcement opportunities [44]. Stretch-attend pose (or stretching) represents an ideal proactive response that allows collecting information about danger in the shape of risk assessment. We thus propose that the expression of defensive behavioral repertoire reflects both threat imminence as well as estimates of agency in APA.

In order to test whether proactive and reactive defensive behaviors share common neural correlates, we artificially modulated levels of arousal in mice trained in inactive and active versions of APA. To do so, we used desipramine which selectively inhibits norepinephrine transporter leading to enhanced extracellular norepinephrine levels [45]. Acute systemic treatment prior to APA caused a drastic enhancement of reactive defensive behaviors (freezing) while sparing proactive defensive behaviors (stretching). Interestingly, desipramine-induced enhancement of freezing behavior in APA was accompanied with a decrease in exploratory behaviors (locomotion, rearing) in male mice only. This effect could be accounted for by the low level of exploratory behavior observed in female mice. The effect observed on exploratory behavior in APA is most likely related to the presence of acute danger, as exploration was spared in its absence. However, as desipramine promotes behavioral habituation, we found a gradual increase in immobility over time. This effect is in line with the observation that norepinephrine expressing terminals in hippocampal CA3 area play a critical role in the acquisition of behavioral habituation to a novel environment [46].

Importantly, mice treated with desipramine not only were able to avoid the footshock zone but also showed greater levels of darting as well as escape vigor as compared to control mice, only when the platform was rotating. This observation echoes the very well-established effect of acute treatment with desipramine in the forced swim test (FST) [47]. In a confined submerged environment, desipramine promotes reactive responding in the shape of enhanced climbing akin to escape behavior [48]. We propose that results obtained in APA and FST reflect a shared mechanism by which increasing extracellular norepinephrine levels enhances reactive defensive responses whenever environment controllability is limited. In APA, desipramine only enhances escape vigor when the platform is rotating, that is, whenever environment controllability is attenuated.

Taken together, our results demonstrate that APA is ideally positioned to study the interaction between danger imminence and estimates of agency on the calibration of proactive and reactive defensive responses. Using different footshock intensities, this paradigm may enable the characterization of mechanistic points of divergence and convergence between sexes. Using different platform revolution speeds, APA may be well suited to perform comparative studies on the neural correlates of proactive and reactive defensive responses. Future work combining APA with the modern neuroscience toolbox will undoubtedly illuminate the neural circuits and molecular mechanisms of individuals response along a behavioral continuum ranging from proactive to reactive responses.

### Limitations

Because we sought to investigate the structure of exploratory and defensive behaviors across danger states in APA, we opted for a supervised definition of behaviors. This strategy contrasts with recent unsupervised approaches that discover behavioral modules in a data-driven manner [49–52]. Future work implementing this comprehensive description of behavioral motifs in APA will undoubtedly provide deeper insights into individual behavioral trajectories in APA. In addition, testing different mouse strains could also unveil differences in the expression of proactive and reactive defensive behaviors.

## METHODS

### Animal Care

10 week-old C57BL/6J mice were purchased from Charles River Laboratories (France). Male (n=64) and female (n=89) mice were housed four per cage (standard sizes according to the European animal welfare guidelines 2010/63/EU) and maintained in a 12 hr light/dark cycle (7:00 a.m. to 7:00 p.m.), in stable conditions of temperature (22°C) and humidity (60%) with ad libitum access to food and water. Age-matched mice (12-13 week-old) were used for behavioral experiments, which took place between 9:00 a.m. and 5:00 p.m. One week prior each experiment, mice were housed individually to prevent aggression and handled daily. Experiments were conducted in accordance with guidelines of the French Agriculture and Forestry ministry for animal care (authorization number/license B34-172-41) and approved by the relevant local and national ethics committees (authorization APAFIS#32853).

### Drugs and treatments

Desipramine (20 mg/kg, Sigma-Aldrich, France) was dissolved in 0.9% NaCl. Desipramine or its vehicle were administered intraperitoneally (i.p.) 30 min prior to the experiment.

### Active place avoidance

The apparatus consisted of a circular (40 cm diameter) electrified stainless steel grid floor that rotated clockwise at a speed of 1 rpm (Delta Technologies Intl / Imetronic, Marcheprime, FR). A clear Plexiglas cylinder (40 cm height) allowed mice to use distal cues present in the room to avoid a 60° stationary shock zone. An overhead camera allowed tracking the position of the mouse in real-time with Poly software (Delta Technologies Intl / Imetronic, Marcheprime, FR). Each entry into the shock zone resulted in a brief constant current footshock (500 ms, 60 Hz, 0.1 to 0.3 mA) that was scrambled across pairs of rods. Failure to exit the shock zone resulted in an additional shock (intershock interval 1.5 s). The platform was divided in six 60° zones and the number of entries in each zone was computed with Poly File software (Delta Technologies Intl / Imetronic, Marcheprime, FR).

### General procedure

Mice were habituated to handling everyday one week prior to the experiment. Mice were brought to the behavioral room at least 30 min prior to the beginning of the experiment. The behavioral room was kept in stable conditions of temperature (22°C) and humidity (60%). On day 1, mice were allowed to explore freely the rotating platform in the absence of shock for 30 min (Pretraining). On days 2 and 3, the shock was turned on and mice were trained to avoid the stationary shock zone defined by cues in the environment for 30 min (Training). On days 4 and 5, the shock was turned off and mice were free to enter the previous shock zone for 30 min (Extinction). Each pretraining, training or extinction session took place once daily for 30 min at roughly the same time (24 hours apart). The apparatus was cleaned with distilled water followed by 70% ethanol between each trial.

### Footshock intensity

In order to test what footshock intensity elicits the most robust active place avoidance, four groups of mice (n=8-16 per group) were trained following the general procedure with either 0 mA, 0.1 mA, 0.2 mA or 0.3 mA during the training phase.

### Visual cues obstruction

In order to test the importance of visual cues to locate the shock zone, one group of mice (n=8) was trained following the general procedure with the exception that proximal cues were covered during the extinction phase (no proximal cues). Another group of mice (n=8) was trained following the general procedure with the exception that an opaque screen was applied onto the clear Plexiglas cylinder to block all visual cues present in the environment during the extinction phase (no visual cues). Results for the

visual cues condition proceed from the footshock intensity experiment (0.3 mA condition).

### Inactive place avoidance

In order to test how platform rotation impacts the learning and extinction of active place avoidance, one group of mice (n=16) was trained following the general procedure (0.3 mA) with the exception that the platform remained immobile during pretraining, training and extinction phases. Results for the active condition proceed from the footshock intensity experiment with rotating platform (0.3 mA condition).

### Desipramine in inactive and active place avoidance

In order to test how platform rotation and desipramine treatment impact active place avoidance, two groups of mice (n=10 per group) were trained in the inactive version of the task with the exception that the platform was turned on during the late phase of training on day 3. One group was treated with desipramine 30 min prior to days 2 and 3 and a control group was treated with vehicle.

### Desipramine in open field and contextual fear conditioning

In order to test the effects of desipramine on spontaneous exploration and contextual fear learning and retrieval, two groups of mice (n=10 per group) were treated with either desipramine or vehicle 30 min prior to exposure to the open field (day 1), contextual fear conditioning (day 2) and retrieval (day 3). The 30 min open field session was performed in the same apparatus as APA with the exception that the platform was immobile and no shock was delivered.

### Contextual fear conditioning

Contextual fear conditioning was performed in the same conditions as APA with the exception that an opaque white chamber was placed onto the electrified grid. The conditioning chambers (20 x 20 x 36 cm) consisted of 4 white walls without ceiling. Freezing behavior was recorded with an overhead camera allowing tracking the position of the mouse in real-time with Poly software (Delta Technologies Intl / Imetronic, Marcheprime, FR). The chamber was cleaned with distilled water followed by 70% ethanol between each trial. The contextual fear conditioning protocol consisted in three 2 s footshocks of 0.3 mA which were delivered every 180 s after placement of the mouse in the training context. The mouse was taken out 20 s after termination of the last footshock. The next day, the animals were briefly re-exposed to the same context (180 s). No footshocks were delivered during the test session. Freezing behavior was measured over 180 s bins preceding each foothsock. Escape vigor was measured at the onset of each footshock over 1 s. Both freezing and escape vigor were analyzed with custom Python scripts that are available online (GitHub).

### Video Recordings

Grayscale 20 fps 16 GB .avi files were downsampled with FormatFactory (v4.5.5) media processing software as 250 MB .avi files. These files were subsequently used for markerless pose estimation analysis.

### DeepLabCut (DLC)

DeepLabCut (v2.3.9) was used to track 10 tracklets (nose, left ear, right ear, neck, left side, right side, center, left hip, right hip, tail). The networks for different tests were trained using 10–20 frames from multiple randomly selected videos for 100,000 iterations. The data generated by DeepLabCut were processed using custom Python scripts that are available online (GitHub).

### Post-hoc identification of arena center and radius

X and Y coordinates of DLC-tracking data were imported into Python (v3.9.7) and processed with custom scripts (GitHub). To identify the center of arena for each experiment, dataframes resulting from each experimental subject obtained on the same day were concatenated. The neck coordinates were plotted and a custom-made graphical user interface was used to define center_x, center_y as well as the radius of the arena surrounding the concatenated path of all subjects (rearing events were clearly distinguishable and not included). A CenterRadius dataframe was generated for each training day that included the values for center_x, center_y, arena_radius+40 (40 pixels wider than the actual radius in order to discard value points beyond the arena), neck_radius (10% wider than the arena radius and used to subsequently measure supported rearing events). Values were then averaged if no discrepancy was encountered across training days (ruling out potential displacement of the platform across sessions).

### Rotation correction of DLC coordinates

The CenterRadius dataframe as well as X and Y coordinates of DLC-tracking data were imported into Python (v 3.9.7) and processed with custom scripts (GitHub). The distance of all tracklets was calculated from the center and body parts beyond the arena were replaced with values from the previous frame. The path of each tracklet was smoothed using a window size of 6 frames. To convert room frame data into arena frame data, 6 degree/sec platform rotation was corrected. Cartesian coordinates were converted into polar coordinates for all tracklets. Theta values were corrected by adding a correction factor of 0.005241667 radian per frame (50 ms). Corrected polar coordinates were then re-expressed as Cartesian coordinates. No correction was applied when the platform was stationary in the inactive place avoidance. To obtain stable data, centroids for head, chest and back were calculated with nose, left ear, right ear and neck tracklets (head), neck, left side, right side and center (chest), center, left hip, right hip and tail (back). Subsequent data analysis was then performed across head, chest and back centroids, unless specified otherwise.

### Analysis of individual behaviors with DLC coordinates

Total locomotion was obtained from chest centroid and was expressed in meters per 30 min. Rearing events reflected the number of times the head centroid crossed a boundary corresponding to 110% arena radius. Center time corresponded to the percent time all centroids spent in the center of the arena (<40% total arena radius). Freezing was determined based on centroids maximal speed <1 cm/sec (rotating platform) or <0.5 cm/sec (stationary platform) and bout duration >2 sec. Stretching events were determined when the speed of the back centroid was <3 cm/sec and the distance between the head and back centroids was >120% the median distance (calculated over 30 min) for > 1 second. Darting events were measured when the speed of the chest centroid was >40 cm/sec anytime but within 1 second following footshocks. Escape vigor was calculated as the median speed of all three centroids for footshock events within 1 second after footshocks.

### Classifiers

Experimental group membership was predicted from behavioral motifs usage (Fig. 3C, 5C), eight features (locomotion, rearing, time in the center, footshock entries, stretching, freezing, darting, and escape vigor) were standardized (z-score). Groups of equal size were classified using a random forest classifier (500 trees) with 5-fold cross-validation (individual observations). Model performance was quantified using accuracy, confusion matrices, and precision/recall metrics. Significance of classification accuracy was assessed via permutation testing, in which group labels were randomly shuffled (500-1000 permutations) to generate a null distribution.

### Blinding

During testing, investigators were not blind to conditions. However, all behavioral data was analyzed automatically using Poly File software (Delta Technologies Intl / Imetronic, Marcheprime, FR) or DeepLabCut (v2.3.9).

### Behavioral quantifications

Quantifications were performed with Python 3.9 (Python Software Foundation) using the following modules: Pandas [53], Matplotlib [54], Numpy [55], Scikit-learn [56], Scipy [57], Plotly [58], Seaborn [59], Statsmodels [60].

### Statistical analysis

No statistical methods were used to pre-determine sample sizes but our sample sizes are similar to those reported in previous publications [61].. Statistical analysis was carried out using GraphPad Prism v10.6 software. Data distribution was assumed to be normal but this was not formally tested unless specified otherwise. Specific details of statistical test, statistics, number of samples and p-values are described in the figure legends and supplementary Table 1. Null hypothesis was rejected with α < 0.05.

### Data exclusion

One mouse was excluded from the inactive condition (Fig. 4) as it was inadvertently placed on the former shock zone during extinction, which dramatically accelerated the extinction rate.

## Data availability

All data generated in this study are available from the lead contact without restriction. Further information and request for original data should be directed to and will be fulfilled by Antoine Besnard.

## Acknowledgements

We thank Drs. Amaury François and Felix Leroy for their valuable comments on the manuscript. We thank iExplore animal facility at the Institute of Functional Genomics, for their help in the maintenance of mouse colonies. We thank Aquineuro for their help in setting up DeepLabCut and Delta Technologies Intl / Imetronic for their help setting up the active place avoidance paradigm. A.B. acknowledges support from the French National Research Agency (ANR-21-CE37-0015), Brain and Behavior Research Foundation NARSAD Young Investigator award (30629), Fondation Fyssen and Fondation pour la Recherche sur le Cerveau.

## Author contributions

A.C., C.Q., S.B., T.V. and A.B. performed behavioral experiments. A.C. and A.B. designed experiments, wrote python scripts and analyzed data. J.N. provided scientific input on automated mouse behavior analysis. F.B. and E.V. contributed to data interpretation and provided scientific input. A.B. conceived and supervised all aspects of the project. A.C. and A.B. wrote the manuscript with assistance from all other authors.

## Declaration of interests

The authors declare no competing financial and non-financial interests.

---

**Figure S1.**
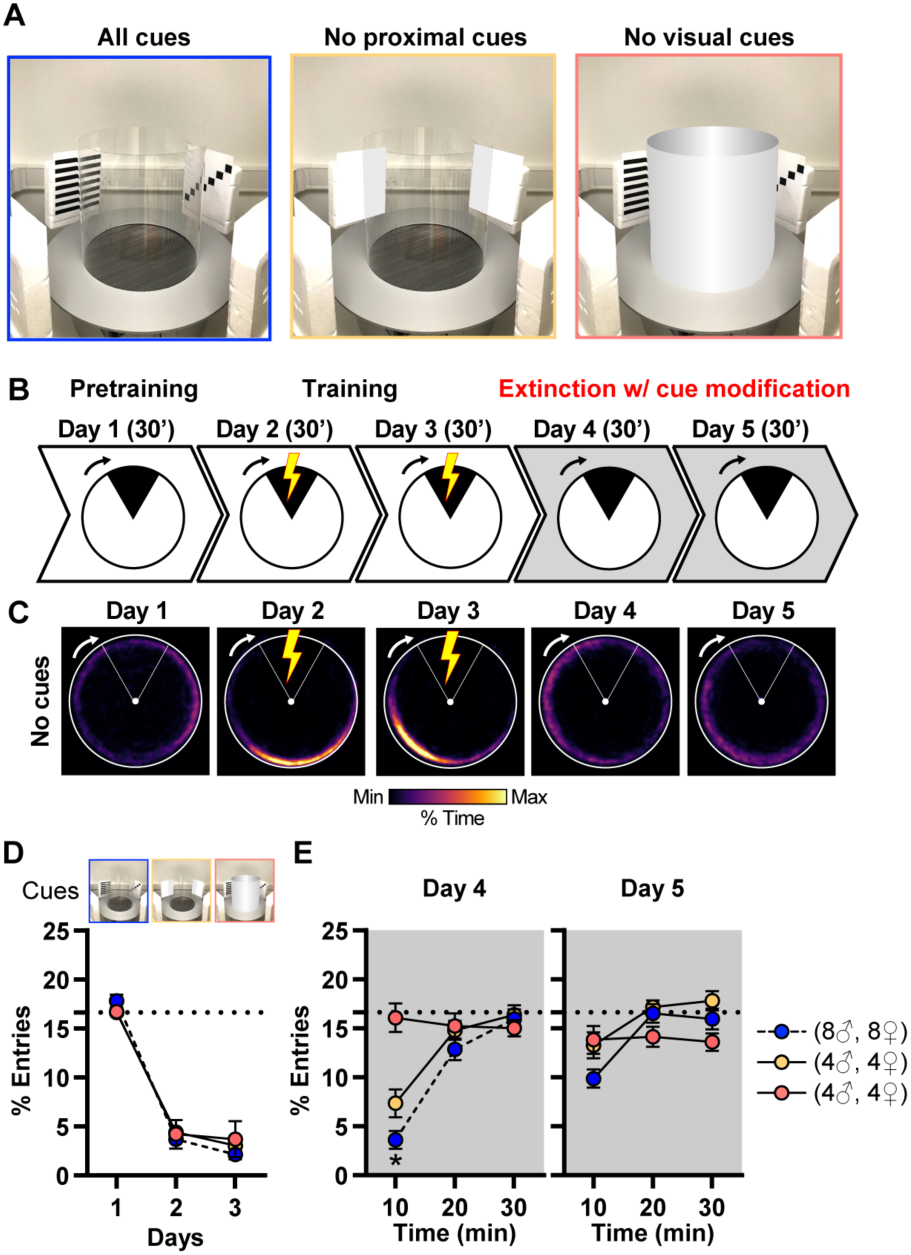
Distal but not proximal cues are necessary for APA (related to Figure 1). **A)** Schematic illustrating the partial (no proximal cues) or complete obstruction of visual cues. **B)** APA consisted of 1 day of pretraining, 2 days of training and 2 days of extinction. Visual cues were altered during the extinction phase on days 4 and 5 (gray area). **C)** Time-in-location heat maps for mice trained with 0.3 mA footshocks and whose visual cues were omitted during extinction training on days 4 and 5. Note the absence of avoidance during both sessions on days 4 and 5. **D)** No difference in the acquisition of APA in the presence of visual cues. **E)** Removal of visual cues blunted the avoidance of the shock zone during the extinction phase. Mice undergoing extinction in the presence of visual cues were the same as the group presented in Fig. 1 (0.3 mA) and was only used for direct comparison. Data (means ± SEM; n= 16, 8, 8 mice per group) were analyzed using mixed factor two-way ANOVA (repeated measure over time) followed by Bonferroni multiple comparisons post-hoc test (detailed in Supplementary Table 1), *p < 0.05, cues versus no cues.

**Figure S2.**
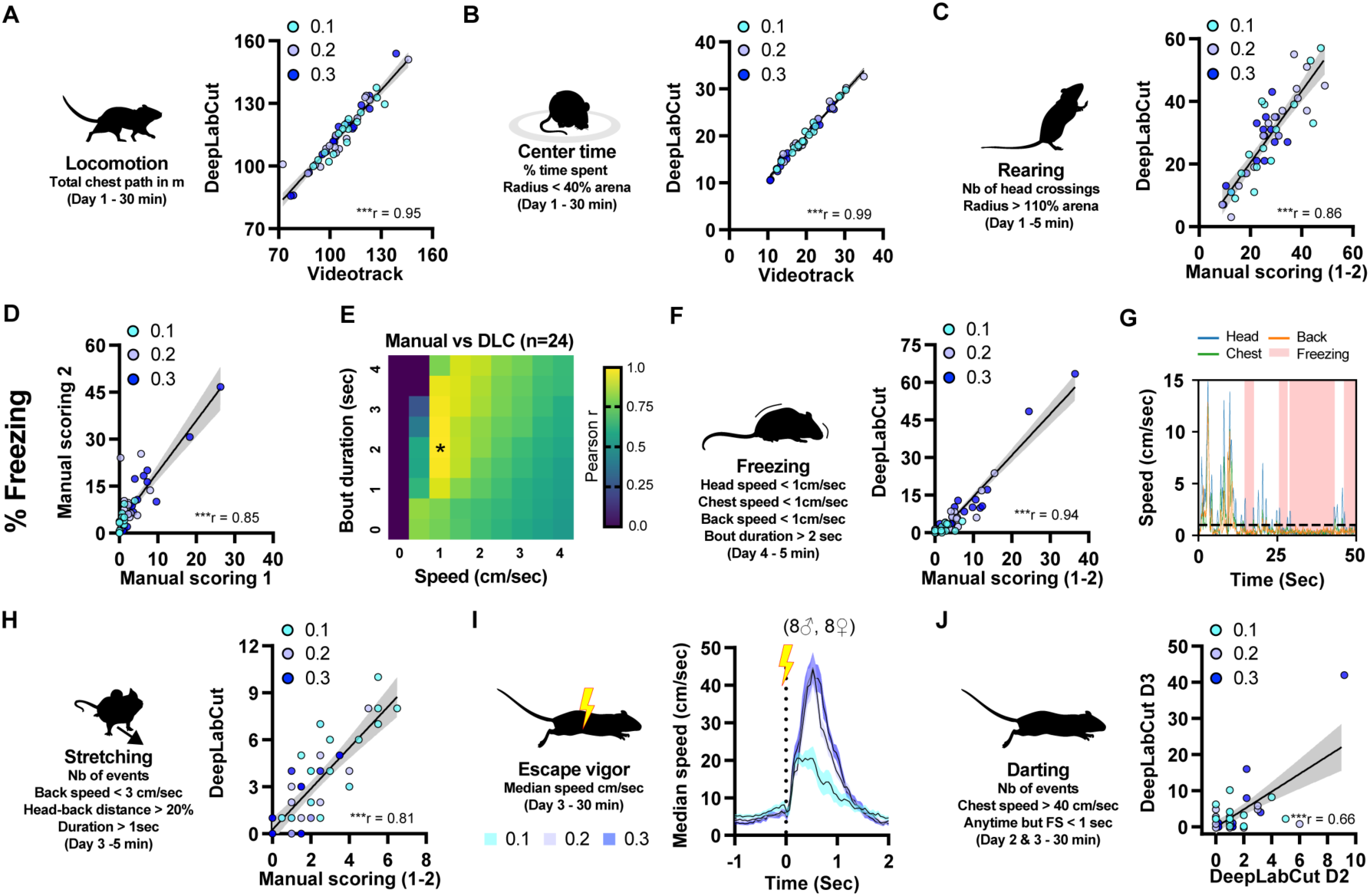
Sex specific differences depend on footshock intensity in APA (related to Figure 2). **A-C)** Total locomotion was obtained from the chest centroid and was expressed in meters per 30 min (A), center time corresponded to the percent time all centroids spent in the center of the arena (<40% total arena radius) (B), rearing events reflected the number of times the head centroid crossed a boundary corresponding to 110% arena radius (C). All DLC measurements reached videotrack or manual scoring accuracy. **D)** Freezing behavior was measured during the first 5 minutes of extinction training on day 4 by two blinded experimenters. Manual measurements were significantly correlated; both measures were thus averaged for subsequent analyses. **E-G)** 24 mice were randomly chosen in order to compare manual scoring (average of both manual scores) with DLC analyses. Freezing was determined based on two criteria, centroids maximal speed and freezing bout duration. Manual scorings were systematically compared with different DLC measurements of varied centroids threshold speed (from 0 to 4 cm/sec) and freezing bout duration (from 0 to 4 sec) (E). DLC and manual scoring reached most significant correlations with a threshold speed of <1 cm/sec (for each centroid) and a bout duration >2 sec (E, F). These detection parameters prevented spurious freezing detection when head centroid was in motion (G). **H)** Stretching events were annotated when the speed of the back centroid was below 3 cm/sec and the distance between the head and back centroids was greater than 120% the median distance (calculated over 30 min) for more than 1 second. **I)** Escape vigor was calculated as the median speed of all three centroids for footshock events aligned from 1 second prior to 2 seconds post-footshocks. **J)** Darting events were measured when the speed of the chest centroid was above 40 cm/sec anytime but within 1 second following footshocks. Darting events were consistent across training sessions.

**Figure S3.**
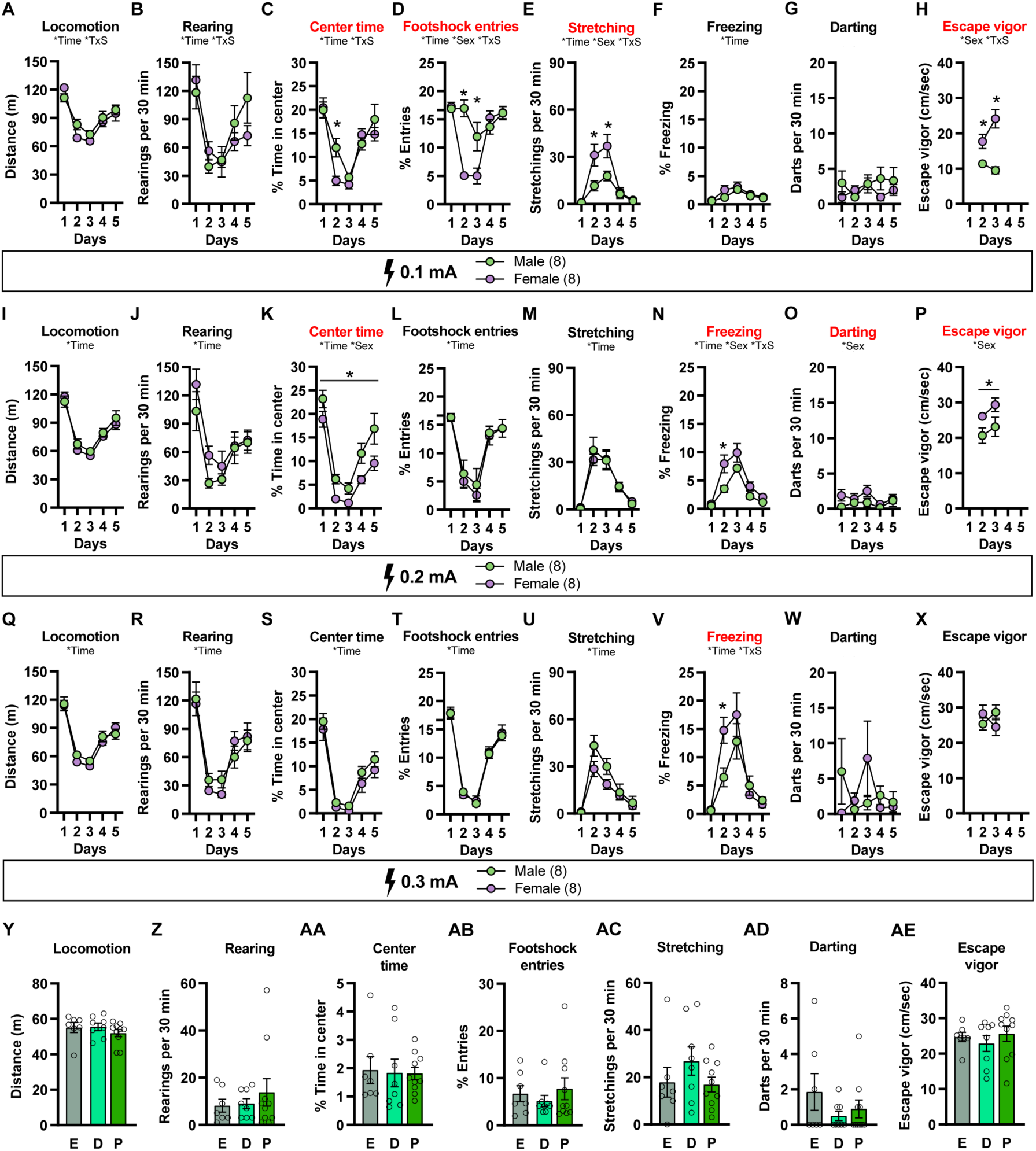
Sex specific differences depend on footshock intensity in APA (related to Figure 3). **A-H)** Pose estimation analyses of locomotion (A), rearing (B), percent time in center (C), footshock entries (D), stretching (E), freezing (F), darting (G) and escape vigor (H) across the 5 days protocol in male and female mice trained with 0.1 mA footshock. Red titles denote significant differences for clarity. Main effect of time or sex as well as interaction (TxS) are indicated for each comparison. **I-X)** Same as A-H except that data were displayed for male and female mice trained with 0.2 mA (I-P) and 0.3 mA (Q-X) footshocks. Red titles denote significant differences for clarity. Data (means ± SEM; n= 8, 8 mice per group) were analyzed using mixed effect model (repeated measure over time) followed by Bonferroni multiple comparisons post-hoc test (detailed in Supplementary Table 1), *p < 0.05, male versus female. **Y-AE)** Pose estimation analyses of locomotion (Y), rearing (Z), percent time in center (AA), footshock entries (AB), stretching (AC), darting (AD) and escape vigor (AE) across mice at different stages of the estrous cycle. Data (means ± SEM; n= 7, 8, 10 mice per group) were analyzed using ordinary one-way ANOVA (detailed in Supplementary Table 1).

**Figure S4.**
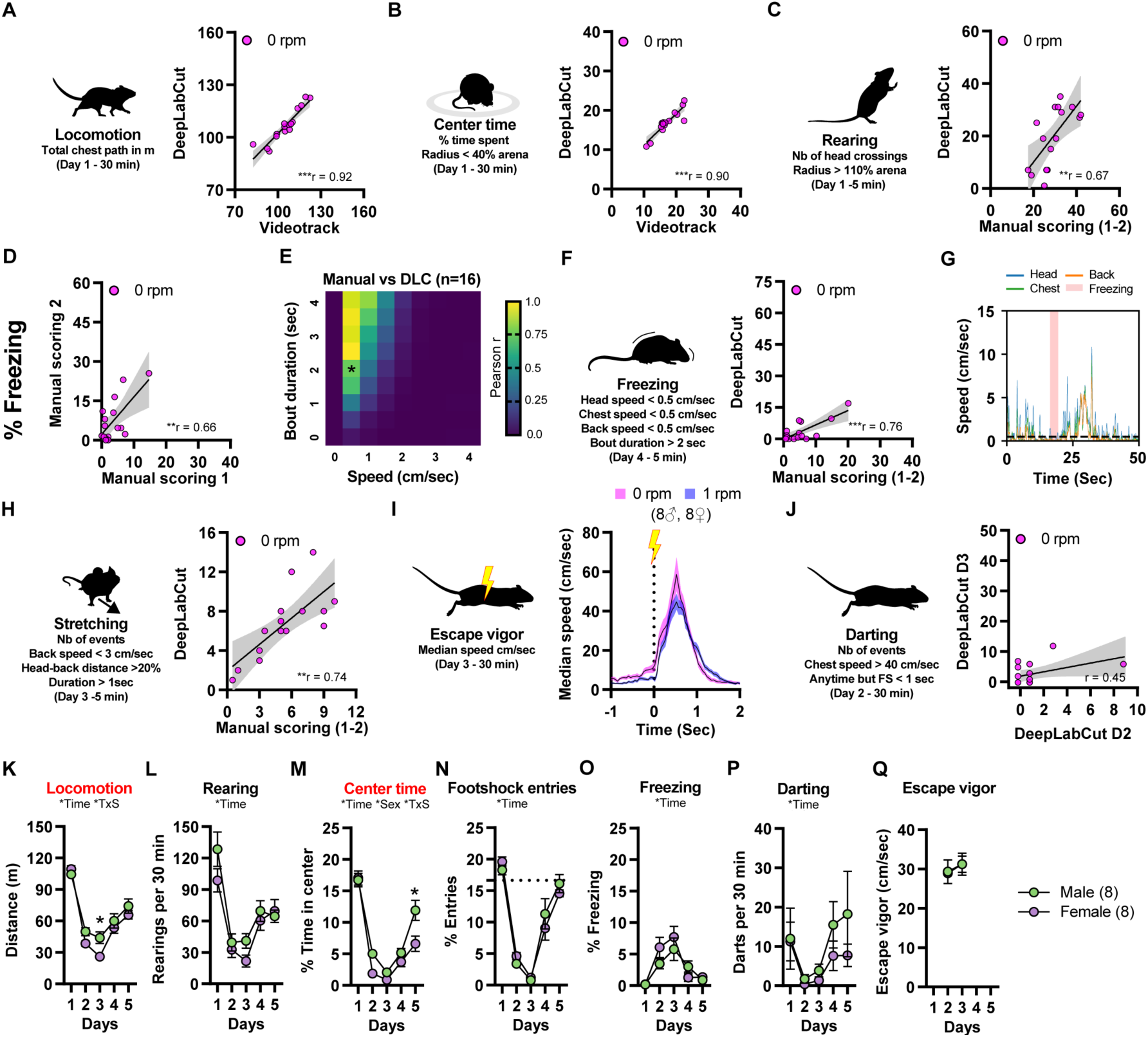
Pose estimation analysis in inactive place avoidance (related to Figure 4). **A-C)** Total locomotion (A), percent time in center (B) and rearing events (C) were analyzed in the inactive version as previously described in the active version, except that no motion correction was performed. All DLC measurements reached videotrack or manual scoring accuracy. **D-G)** Freezing behavior was analyzed in the inactive version as previously described in the active version, except that no motion correction was performed. Note that the speed threshold is set to 0.5 cm/sec in the inactive version (E), which matches the 1 cm/sec in the active version owing to a subsiding jitter when performing motion correction. **H-J)** Stretching events (H), escape vigor (I) and darting events (J) were annotated in the inactive version as previously described in the active version, except that no motion correction was performed. **K-Q)** Pose estimation analyses of locomotion (K), rearing (L), percent time in center (M), footshock entries (N), freezing (O), darting (P) and escape vigor (Q) across the 5 days protocol in male and female mice trained in inactive place avoidance. Red titles denote significant differences for clarity. Data (means ± SEM; n= 8, 8 mice per group) were analyzed using mixed-effect model followed by Bonferroni multiple comparisons post-hoc test (detailed in Supplementary Table 1), *p < 0.05, male versus female. Main effect of time or sex as well as interaction (TxS) are indicated for each comparison.

**Figure S5.**
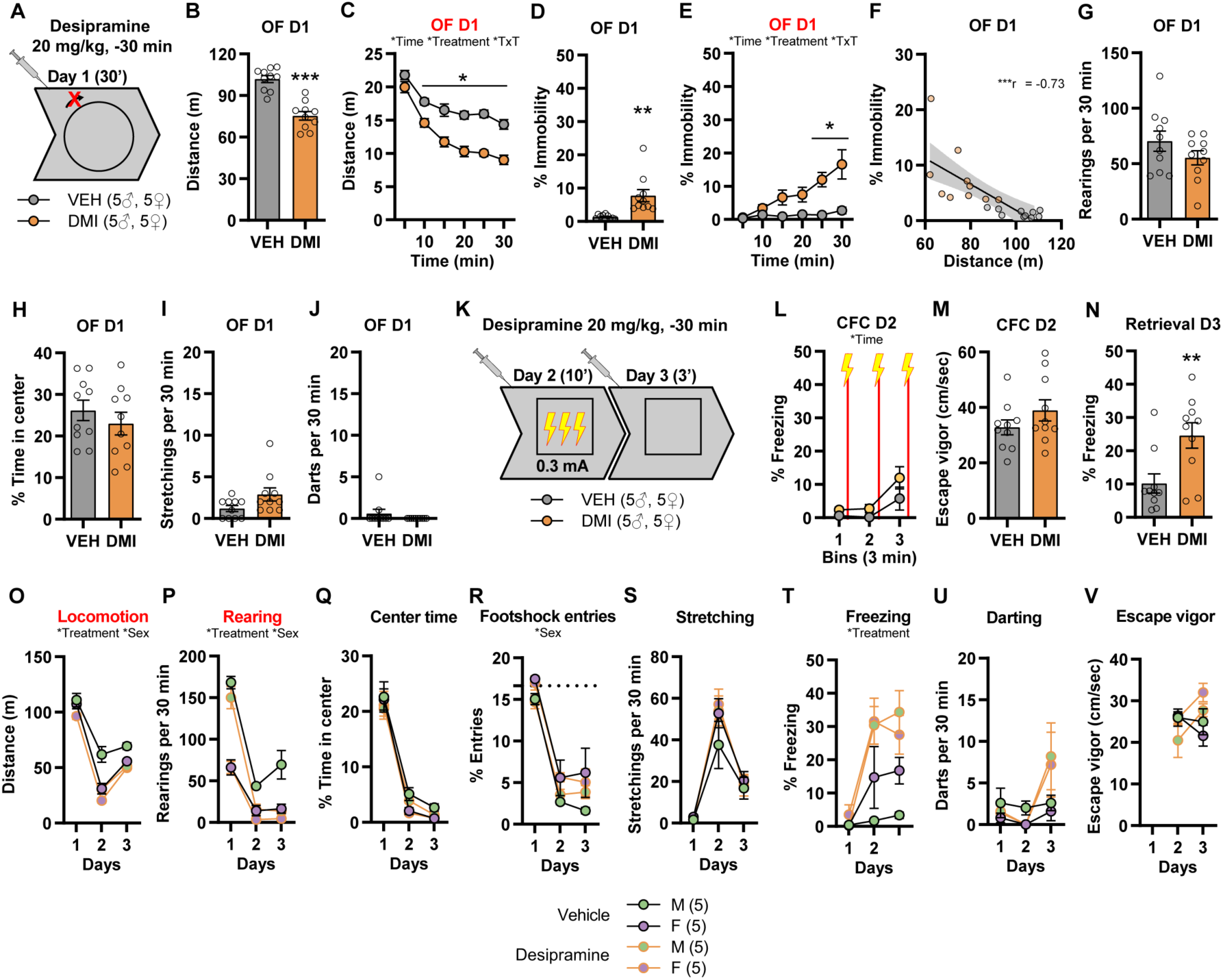
Enhancing norepinephrine transmission promotes behavioral habituation in an open field and contextual fear memory retrieval (related to Figure 5). A-J) Acute injection with desipramine prior to open field exposure (A), decreases locomotor behavior (B), in a time-dependent manner (C), increases immobility (D), in a time-dependent manner (E). Immobility is inversely correlated with locomotion in mice treated with desipramine or vehicle (F). Acute desipramine injection has no effect on rearing (G), percent time spent in center (H), stretching (I) and darting (J) in the open field. Main effect of time or treatment as well as interaction (TxT) are indicated for each comparison. K-N) Acute injection with desipramine prior to contextual fear conditioning on subsequent days 2 and 3 (K), has no effect on contextual fear conditioning (CFC) with 0.3 mA electric footshocks (L), escape vigor elicited by footshocks during CFC (M), but enhances freezing behavior during a brief retrieval session on day 3 (N). Data (means ± SEM; n= 10, 10 mice per group) were analyzed using mixed-effect model followed by Bonferroni multiple comparisons post-hoc test (detailed in Supplementary Table 1), *p < 0.05, desipramine versus vehicle. O-V) Pose estimation analyses of locomotion (O), rearing (P), percent time in center (Q), footshock entries (R), stretching (S), freezing (T), darting (U) and escape vigor (V) across the 3 days protocol in male and female mice treated with desipramine or vehicle. Red titles denote significant differences for clarity. Data (means ± SEM; n= 5 mice per group) were analyzed using mixed factor three-way ANOVA (repeated measure over time) (detailed in Supplementary Table 1). Main effect of treatment or sex are indicated for each comparison

## Notes

### Competing Interest Statement

The authors have declared no competing interest.

